# HIV-1 latency reversing agents converge on phosphoregulation of nuclear protein complexes

**DOI:** 10.1101/2025.11.13.688376

**Authors:** Jocelyn J. Ni, Prashant Kaushal, Shipra Sharma, Sara K. Makanani, Yennifer Delgado, Sophie F. Blanc, Charles Ochieng Olwal, Immy A. Ashley, Declan M. Winters, Vivian Yang, Callie N. Phung, Kareem Alba, Evan Wynn, Jeffrey R. Johnson, Judd F. Hultquist, Oliver I. Fregoso, Mehdi Bouhaddou

## Abstract

Despite the success of antiretroviral therapy (ART), HIV-1 persists in latently infected cells, posing a central barrier to a cure. “Shock-and-kill” strategies using latency-reversing agents (LRAs) have shown some promise in reactivating viral gene expression *ex vivo*, but have yielded little clinical efficacy, underscoring the need for deeper insight into the molecular mechanisms that govern reactivation. Here, we performed deep quantitative phosphoproteomics of J-Lat 10.6 cells treated with diverse LRAs: SAHA, PMA, or prostratin. We identified 48,476 confidently localized phosphorylation sites mapping to 6,672 proteins, with SAHA inducing the most extensive changes. Regulated phosphoproteins were enriched in chromatin organization, transcription, RNA processing, nuclear transport, and cytoskeletal remodeling. Although LRAs regulated overlapping pathways, they elicited divergent kinase activities and site-specific phosphorylation patterns. A reproducible core of 3,502 phosphorylation sites on 1,432 proteins mapped to 39 nuclear protein complexes, including the spliceosome, Mediator, NF-κB, and RNA polymerase II. Remarkably, 20 protein complexes were phosphoregulated by all three LRAs, but at distinct sites, revealing convergence on shared nuclear machinery through distinct mechanisms. This study provides a comprehensive map of protein complex phosphorylation remodeling during HIV-1 reactivation and highlights signaling mechanisms that could guide the rational design of next-generation LRAs with improved efficacy and reduced toxicity.

## INTRODUCTION

Despite the success of antiretroviral therapy (ART) in suppressing viral replication, HIV persists as a latent reservoir within long-lived cell types, including memory CD4⁺ T cells, macrophages, dendritic cells, monocytes, and tissue-resident cells, such as microglia and astrocytes in the central nervous system^1,2^. Latent proviruses are replication competent, but transcriptionally or post-transcriptionally inhibited and subject to spontaneous reactivation. This reservoir poses a major barrier to an HIV-1 cure as it facilitates rapid viral rebound upon interruption of ART, emphasizing the need for strategies that eliminate latent proviruses^3–5^.

One proposed curative strategy, termed “shock and kill,” employs latency-reversing agents (LRAs) to induce reactivation of latent HIV-1 and thereby promote recognition and clearance by the adaptive immune system, either alone or in combination with emerging modalities such as CAR T cell therapy^6,7^. Multiple classes of LRAs have been identified, including histone deacetylase inhibitors (HDACi), such as suberoylanilide hydroxamic acid (SAHA), and protein kinase C (PKC) agonists, including phorbol 12-myristate 13-acetate (PMA) and prostratin. SAHA (also known as vorinostat) is an FDA-approved drug used to treat cutaneous T cell lymphoma^8,9^, whereas PMA and prostratin are commonly used experimental LRAs that activate PKC signaling^10,11^.

Thus far, LRAs have shown limited success in achieving HIV-1 cure outcomes. For instance, no LRA has been able to fully reactivate the latent HIV-1 reservoir, particularly in resting CD4⁺ T cells. The most potent LRAs, PKC agonists, induce strong global T cell activation and inflammation, making them too toxic for therapeutic use^12^. Conversely, agents that are more selective, such as HDAC and bromodomain inhibitors, are better tolerated but only weakly reactivate latent proviruses^6,7^. Consistent with these limitations, clinical trials have demonstrated that none of the LRAs tested to date have significantly reduced the size of the latent reservoir or delayed viral rebound following ART interruption^13–19^. These limitations underscore the ongoing need to identify new LRAs that are both potent and safe.

A major challenge in developing improved LRAs is the incomplete understanding of the molecular mechanisms that govern HIV-1 latency and reactivation. Latency is maintained through multilayered transcriptional, post-transcriptional, and post-translational processes^20–22^. Among these mechanisms, protein phosphorylation and protein-protein interactions are central to signaling cascades that coordinate chromatin remodeling, transcriptional activation, and nuclear transport, processes that ultimately determine whether the provirus remains silent or reactivates^23–25^. For example, P-TEFb, a cyclin-dependent kinase complex composed of CDK9 and cyclin T1, is a key regulator of RNA polymerase II transcriptional elongation and is required for efficient HIV-1 latency reversal^26^. The phosphorylation of CDK9 at Serine 175 (S175) enhances P-TEFb complex activity and promotes its recruitment to sites of nascent viral transcription, facilitating transcriptional elongation of viral RNA^27^. These phosphorylation-dependent mechanisms are thought to be essential for initiating and sustaining reactivation; however, a comprehensive, systems-level map of how LRAs remodel phosphorylation and protein complexes has yet to be defined.

We reasoned that systematic characterization of protein complex phosphorylation during LRA-induced reactivation would uncover key host regulatory pathways and potential therapeutic targets. Insights gained could enable the rational design of agents that modulate protein-protein interactions to achieve potent reactivation with reduced toxicity. To test this, we performed a global phosphoproteomics analysis using the J-Lat 10.6 model of HIV-1 latency^28^, profiling cellular responses to three LRAs—SAHA, PMA, and prostratin—in quadruplicates, across two independent experiments at 24 hours post LRA administration. Through multilevel statistical filtering, quantitative comparison, and network analysis, we identified shared pathway-level regulation that occurred through distinct mechanisms. We observed convergent phosphoregulation of a set of nuclear protein complexes, revealing that distinct LRAs regulate the same complexes through divergent mechanisms. Our findings establish a systems-level framework for understanding the host signaling architecture underlying latency reversal and for guiding the development of next-generation LRAs with improved specificity, efficacy, and safety.

## RESULTS

### LRAs induce marked phosphoproteome remodeling

To understand how LRAs influence phosphorylation signaling during HIV-1 reactivation, we performed global phosphoproteomics of J-Lat 10.6 cells^28^, a T cell model of latent HIV with a GFP reporter, exposed to different LRAs. Cells were treated for 24 hours with 0.01% DMSO (control) or one of three LRAs: 10 μM SAHA, 5 ng/ml PMA, or 5 μM prostratin. Each treatment was performed in quadruplicate and was repeated in two independent experiments. After 24 hours of treatment, cells were collected for flow cytometry, mass spectrometry abundance proteomics, and phosphoproteomics. Ten percent of protein was used for global abundance analysis, and 90% for phosphopeptide enrichment, per replicate. Data were processed in Spectronaut and filtered for phosphorylation sites with more than 90% localization probability for high confidence assignment (Fig. 1A).

**Figure 1.**
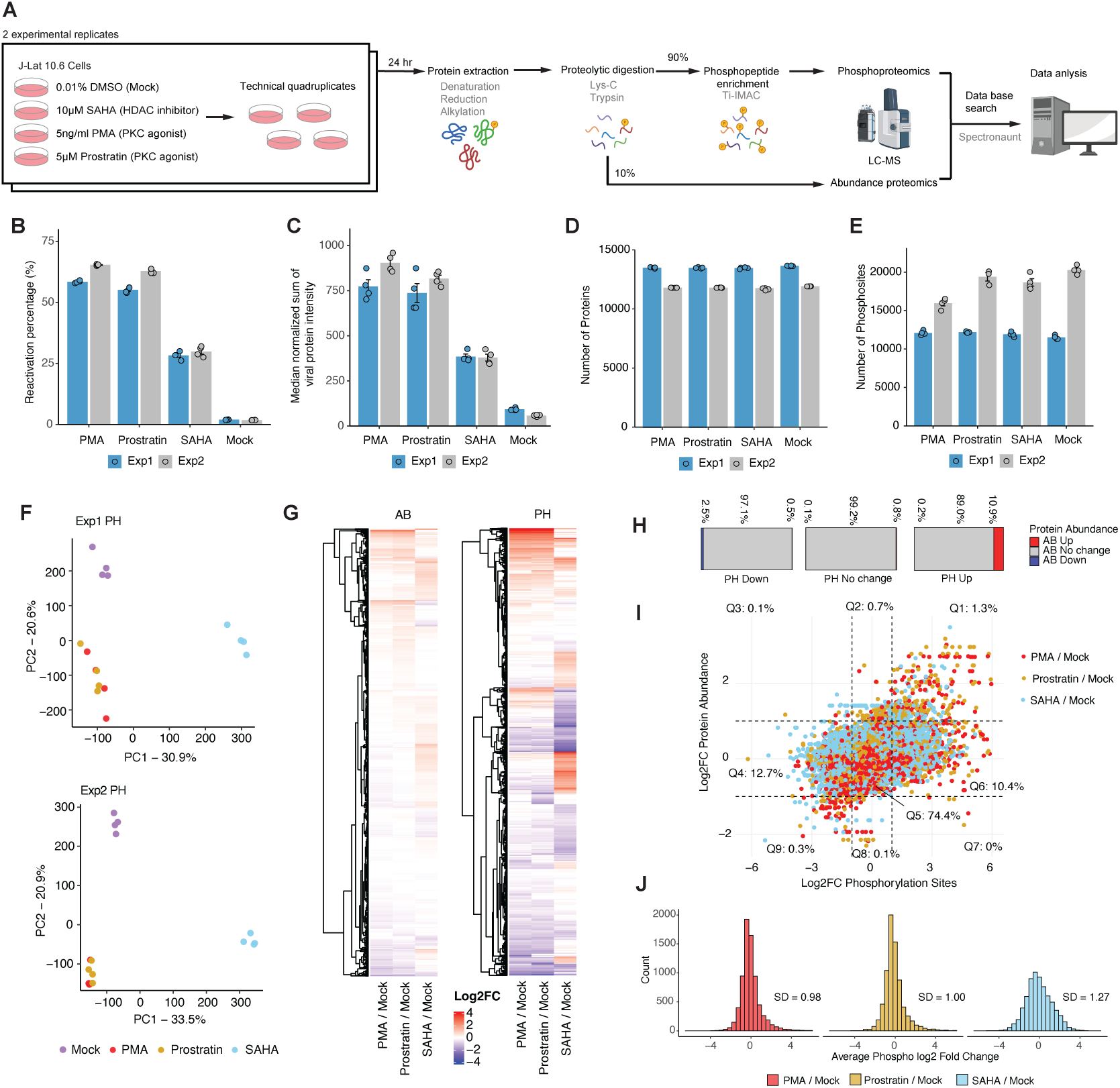
Latency reversal agents induce global remodeling of the phosphoproteome. (A) Experimental workflow. J-Lat 10.6 cells were treated with DMSO, SAHA, PMA, or prostratin for 24 hours in biological quadruplicates across two independent experiments. (B) Flow cytometry quantification of GFP-positive (*i.e.*, reactivated) cells shows reactivation efficiencies for each LRA, with bars depicting the mean of quadruplicates per experiment. (C) Count of viral proteins detected by mass spectrometry reflects reactivation levels, with bars depicting the mean of cross-run normalized (global median) sum of viral protein intensities across quadruplicates, per experiment. (D) Number of proteins identified from the abundance proteomics analysis per LRA treatment and replicate. (E) Number of high confidence (localization probability > 0.9) phosphorylated sites identified from the phosphoproteomics analysis per LRA treatment and replicate. (F) Principal component analysis (PCA) of phosphoproteomic (PH) profiles showing replicate reproducibility and similarities/differences between treatments. (G) Global overview of log₂ fold changes (log₂FCs) in abundance (AB, left) and phosphorylation (PH, right) versus mock, representing the DMSO control, taking average values between Exp1 & 2. (H) For phosphorylation sites downregulated, unchanged, or upregulated in treatment versus DMSO, bar plots depict the fraction with a corresponding downregulation (blue), no change (grey), or upregulation (red) in total protein levels, taking average values between Exp1 & 2. (I) Scatterplot comparing log₂FCs in phosphorylation and protein abundance for all detected phosphorylation sites and proteins. Each dot represents one phosphorylation site or protein, colored by treatment, taking average values between Exp1 & 2. (J) Distribution of phosphorylation log₂FCs for each treatment, showing prostrain and PMA with similar distributions (standard deviation of 0.98 and 1, respectively) and SAHA with a broader range (standard deviation of 1.27).

Flow cytometry analysis of GFP-positive cells 24 hours post-treatment found an average of 62.0%, 59.0%, 29.1%, and 1.9% GFP-positive (*i.e.*, reactivated) cells for PMA, prostratin, SAHA, and DMSO exposure, respectively, in experiment 1 (Exp1) and experiment 2 (Exp2) (Fig. 1B, S1A). GFP-based measures of reactivation by flow cytometry were corroborated by the sum of HIV-1 protein intensities detected by MS abundance proteomics (Fig. 1C). Our phosphoproteomics analysis detected 11,308-20,970 confidently localized (localization probability > 0.9) phosphorylated sites (Fig. 1E) on 3,603-5,221 proteins (Fig. S1C) across replicates and experiments. The abundance proteomics data identified 201,728-236,107 peptides (Fig. S1B) and 11,757-13,606 proteins (Fig. 1D) across replicates and experiments. Principal component analysis (PCA) showed a clear separation based on treatment condition and high reproducibility among replicates for the phosphoproteomics (Fig. 1F, S1D) and abundance data (Fig. S1E). As expected, PMA and prostratin clustered together in the phosphoproteomics dataset, reflecting their similar mechanisms of action (*i.e.*, they are both PKC agonists).

We observed marked phosphoproteome remodeling with high log₂ fold-changes (log₂FC) across conditions, with fewer changes at the level of protein abundance (Fig. 1G). Specifically, of downregulated phosphorylation sites (*i.e.*, log₂FC < -1), only 3% changed in abundance (0.5% up and 2.5% down); of upregulated sites (i.e., log₂FC > 1), 11.1% changed in abundance (10.9% up and 0.2% down); and of unchanged sites (i.e., log₂FC > -1 and log₂FC < 1), 0.9% changed in abundance (0.8% up and 0.1% down) (Fig. 1H, 1I). This analysis highlighted that the majority of changes in phosphorylation were not driven by a change in total protein levels. Relative to DMSO, the distribution of phosphorylation changes was broader for SAHA (standard deviation of 1.27) than for PMA (0.98) or prostratin (1.00; Fig. 1J), suggesting SAHA had a stronger overall impact on the phosphoproteome, which was unexpected given that PMA and prostratin are direct kinase activators.

### LRAs regulate overlapping cellular signaling pathways via distinct mechanisms

We detected similar numbers of differentially abundant proteins (Fig. 2A, left) and phosphosites (Fig. 2A, right) between experiments 1 and 2, with a slight enhancement in experiment 2 (abs[log₂FC] ≥ 1 and q < 0.05). In general, all treatments increased protein abundances, whereas phosphorylation sites showed changes in both directions (Fig. 2A). Since our study was designed to identify shared mechanisms between distinct LRAs, we reasoned that after 24 hours of treatment, production of reactivated viral proteins could drive overlapping pathway modulation reflective of viral protein activity rather than shared pathway regulation by the original stimulus. To assess this, we reperformed our experiment with one LRA of each class, PMA and SAHA, at 2 hours post-treatment. As expected, we did not observe a change in viral protein expression, as evaluated by MS abundance proteomics (Fig. 2B). However, we did observe significant changes in host protein and phosphosite levels (Fig. 2C, 2D); the standard deviation in log₂ fold changes in phosphorylation was 1.33 and 2.33 for PMA and SAHA, respectively (Fig. 2E), similar to what was observed for the 24 hour perturbation (Fig. 1J).

**Figure 2.**
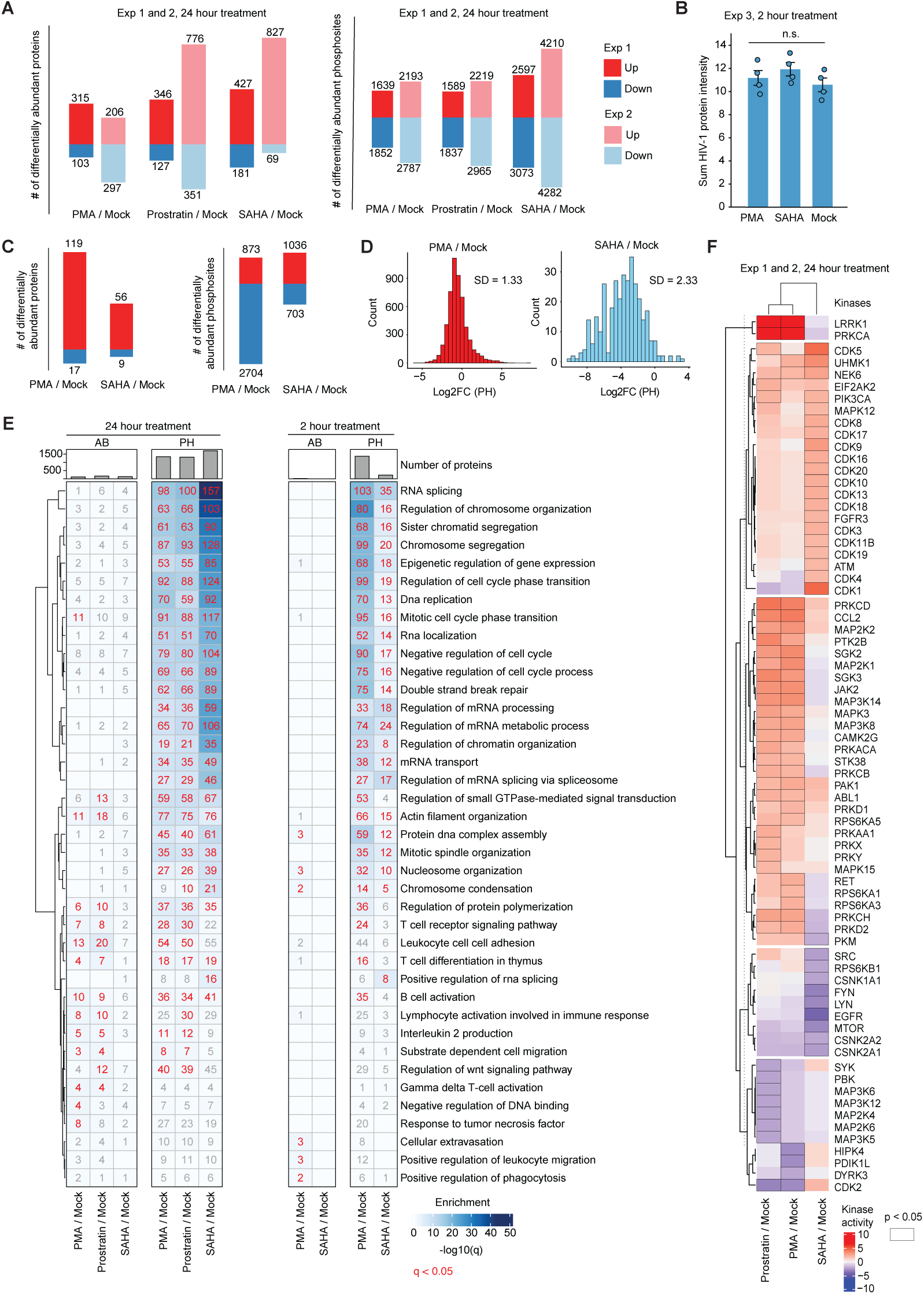
Latency-reversing agents regulate overlapping cellular signaling pathways via distinct mechanisms. (A) Numbers of differentially abundant proteins and phosphosites identified in each condition from unfiltered data across two 24 hour treatment experiments (Exp1 & 2). (B) Sum of cross run-normalized viral protein intensities detected in 2 hour treatment (Exp3), with bars depicting the mean of quadruplicates per experiment. (C) Numbers of differentially abundant proteins and phosphosites identified in each condition from unfiltered data from the 2 hour treatment experiment (Exp3). (D) Distribution of phosphorylation log₂ fold changes across conditions in the 2 hour treatment experiment (Exp3). (E) Gene set overrepresentation analysis (GSOA) based on Gene Ontology Biological Processes (GOBP) on all 2 hour and 24 hour treatment phosphoproteomics and abundance data. (F) Kinase activity analysis on averaged log₂ fold changes across Exp 1 and 2 based on statistical analysis of prior knowledge networks of kinase-substrate interactions (see Methods).

We then performed a gene set overrepresentation analysis (GSOA)^29,30^ on all three datasets, which included the union of proteins or phosphosites differentially regulated in experiment 1 or 2 (24 hour) and all in experiment 3 (2 hour; Fig. 2E). We observed the 24 hour and 2 hour perturbations to reflect highly overlapping pathway-level regulation (Fig. 2E). Protein abundance changes were modest, largely indicating T cell activation and innate immune responses at 24 hours, with minimal changes at 2 hours. Changes in protein phosphorylation were robust and represented diverse pathways, including chromosome organization, epigenetic regulation, DNA replication, DNA damage, RNA splicing, RNA transport, RNA processing, and cytoskeleton organization. Interestingly, the majority of pathways were predominantly nuclear. In general, SAHA showed stronger and more specific enrichment in RNA splicing, chromosome organization, chromosome segregation, and epigenetic regulation of gene expression, as expected given its role in modulating histone deacetylase activity.

Although all three LRAs depicted convergent regulation of cellular pathways, we next asked whether they also overlapped in their regulation of kinase activities. To evaluate this, we performed kinase activity analysis on phosphorylation sites detected consistently across experiments 1 and 2 for the 24 hour time point. Briefly, averaged log₂ fold changes in phosphosite intensities across experiments, between treatment and mock, were used to infer kinase activity states using prior knowledge networks of kinase-substrate interactions^31–34^ (Fig. 2A). We observed that while PMA and prostratin exhibited highly similar kinase activity profiles, SAHA was markedly distinct. As expected, PMA and prostratin robustly activated PRKCA, the catalytic subunit of protein kinase C, and other PKC isoforms (PRKCB, PRKCD, and PRKC), consistent with their roles as PKC agonists. PKC activation was accompanied by strong activation of downstream ERK and JAK-STAT pathways, which indicated broad phosphorylation-driven regulation of stress response programs. Interestingly, SAHA elicited a distinct signaling response at the level of inferred kinase activities compared to PMA and prostratin, predominantly resulting in the activation of cyclin-dependent kinases (CDKs), including CDK1 and CDK4, and the inhibition of EGFR, SRC, and mTOR^35^, leaving ERK signaling unaffected. We concluded that although all three LRAs regulated similar cellular signaling pathways (Fig. 2E), closer examination of dysregulated kinase activities revealed distinct underlying mechanisms of signaling control (Fig. 2F), indicating that distinct LRA classes engage similar pathways through different routes.

### LRAs converge on nuclear protein complexes

We then sought to identify a core set of phosphosites with reproducible changes across experiments. After identifying phosphosites commonly detected in experiments 1 and 2 (24 hour), we retained only those that changed significantly (q < 0.05) and in the same direction. Furthermore, we removed phosphosites on a protein that possessed a corresponding change in total levels to exclude abundance-driven effects (Fig. 3A). The largest reduction occurred when requiring overlap (Step 2) and direction consistency (Step 3) across experiments, while abundance correction removed only a small proportion of sites (Step 4; Fig. 3B).

**Figure 3.**
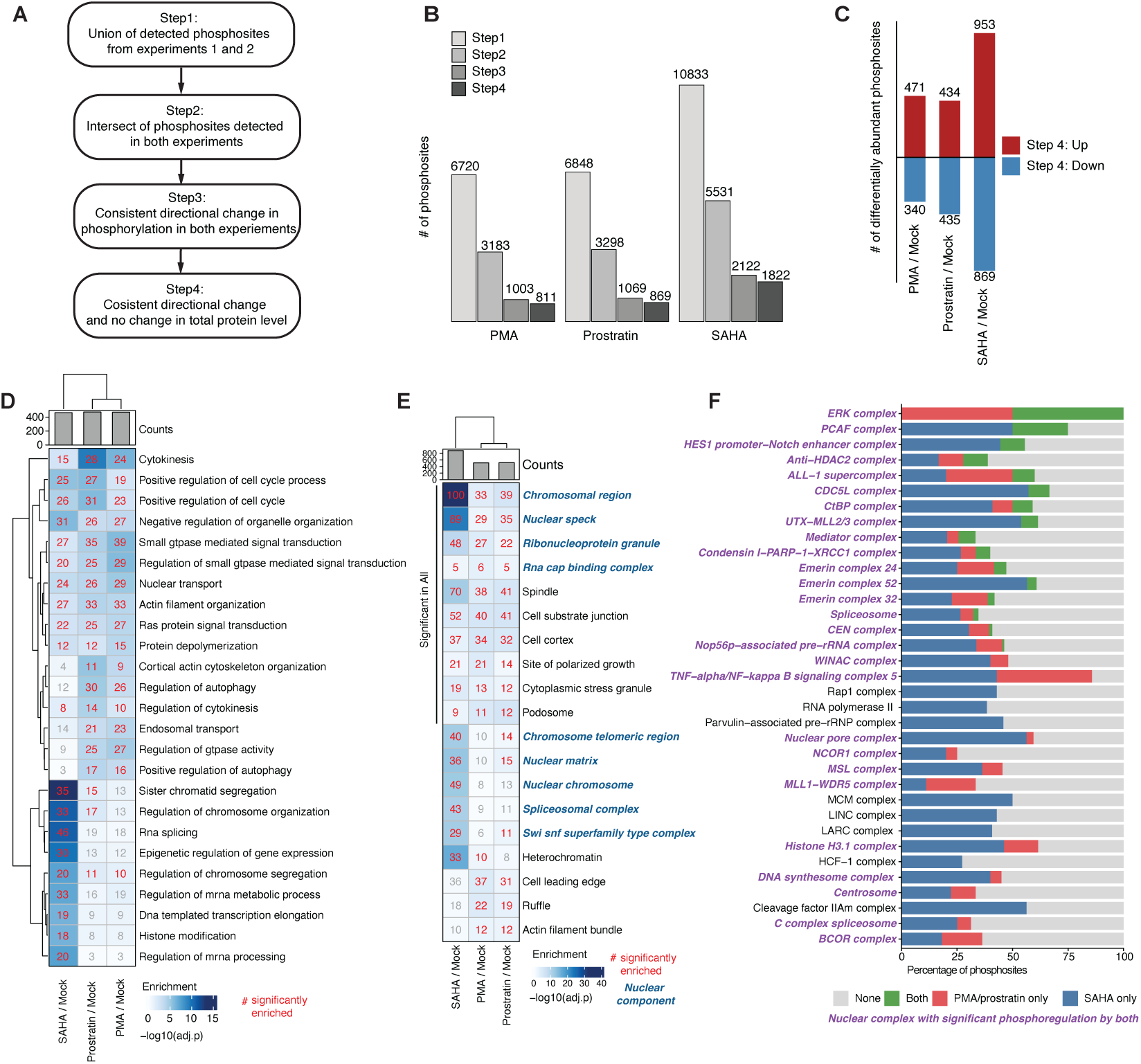
Latency-reversing agents converge on nuclear protein complexes. (A) Flow chart showing multilevel filtering: overlap across experiments (Step 2), direction consistency filtering (Step 3), and abundance correction (Step 4). (B) Number of phosphosites remaining per LRA for each filtering step. (C) Significantly changed phosphosites going up (red) or down (blue) from our final core dataset containing 3,502 abundance-independent significant phosphosites. (D) GSOA on Gene Ontology Biological Pathways (GOBP) of core phosphosites. (E) GSOA on Gene Ontology Cellular Component (GOCC) of core phosphosites. (F) Percentage of significant phosphosites regulated on nuclear protein complexes by SAHA (blue), PMA/prostratin only (red), both (green), none (grey). Complexes with significant phosphoregulation by both LRA classes (*i.e.*, SAHA and PMA/prostratin) are labeled in purple.

After filtering, we found 3,502 phosphosites on 1,432 proteins that changed consistently in both experiments and were abundance-independent, with SAHA contributing the largest number of changing sites (Fig. 3B, 3C). GSOA on these core phosphoregulated proteins uncovered converging pathways, particularly related to cell cycle, nuclear transport, cytoskeleton organization, and GTPase activity (Fig. 3D). SAHA was strongly enriched for chromatin organization, histone modification, and transcriptional regulation (Fig. 3D). Further analysis of the cellular compartments of these core proteins revealed strong enrichment for nuclear regions, including chromatin, nuclear speck, ribonucleoprotein granule, nuclear matrix, and splicing, highlighting convergent regulation in the nucleus (Fig. 3E).

We next used the CORUM database^36^, which curates mammalian protein complexes, to map our core phosphosite set onto nuclear protein complexes; we detected 39 in our dataset. Twenty nine of these complexes were significantly phosphoregulated by both classes of LRAs (*i.e.*, SAHA and PMA or prostratin). Twenty of these complexes were significantly phosphoregulated by all three LRAs (Fig. 3F, bold). Interestingly, different classes of LRAs rarely regulated identical sites; more often, each complex was modulated by different LRAs acting on distinct proteins and/or phosphorylation sites, reinforcing that distinct LRAs converge on similar biological processes through unique signaling mechanisms. Notably, the regulation of a protein complex through different molecular routes may yield comparable functional outcomes, which warrants future investigation.

### Distinct phosphorylation events modulate a core set of nuclear complexes during reactivation

We next constructed a protein-protein interaction network of the 39 nuclear complexes with significant phosphoregulation by at least one LRA from our core (*i.e.*, filtered, Fig. 3C) phosphorylation dataset (Fig. 4). Each node represents a phosphorylated protein, with edges connecting proteins within the same complex. Node rings indicate the LRA condition in which phosphorylation was detected, color indicates the average log_2_-fold changes relative to mock in the two experiments, and phosphorylation sites are labeled within each node segment.

**Figure 4.**
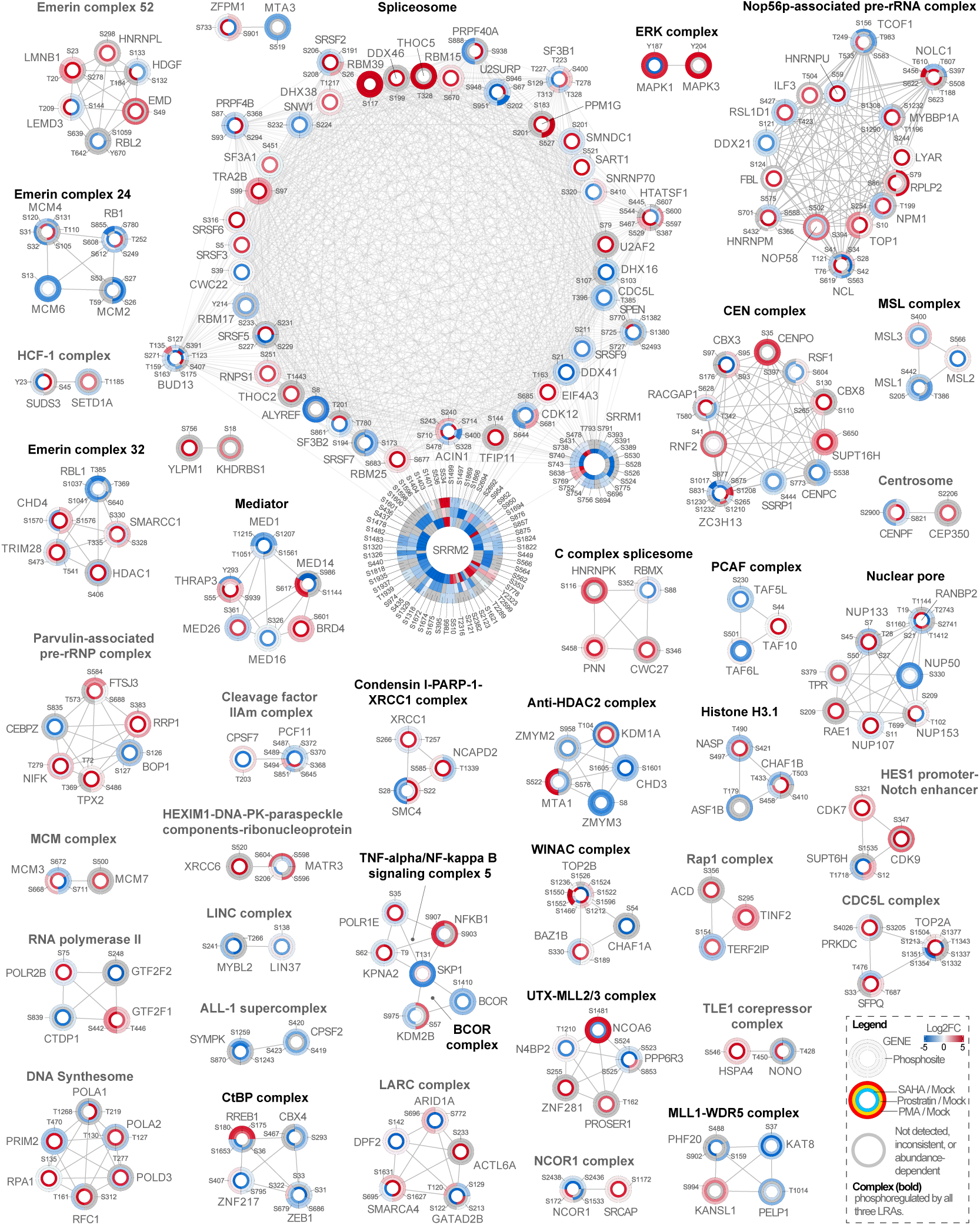
Phosphoregulation of nuclear protein complexes during HIV-1 latency reversal. Protein-protein interaction network of nuclear proteins (circles) containing at least one significant phosphosite in any LRA condition. Each node ring indicates a treatment condition (SAHA: innermost ring; prostratin: middle ring; PMA: outermost ring); node segments are labeled by phosphosite position, and color indicates upregulation (red) and downregulation (blue) of the phosphosite. Bold complex names indicate complexes with significant phosphorylation changes in all three LRAs. Edges connect proteins within the same CORUM-defined complex. n = 2 biologically independent experiments.

The spliceosome was among the most prominently regulated modules in our network, emphasizing the importance of host RNA-processing machinery in HIV reactivation. The most highly phosphorylated proteins included SRRM2 and SRRM1, both serine/arginine-rich splicing factors that coordinate alternative splicing and splice-site selection. SRRM2 has previously been shown to undergo rapid phosphorylation upon HIV-1 entry into CD4⁺ T cells, where it modulates host splicing machinery to facilitate viral replication and release, suggesting a broader regulatory role of SR proteins throughout the viral life cycle^37^. RBM39 and THOC5 depicted prominent increases in phosphorylation, with RBM39 S117 being significantly upregulated in response to all three LRA treatments. Although these proteins have not been directly studied in the context of HIV-1 infection, they are recognized regulators of transcription, RNA processing, and nuclear export and are being explored in cancer as prognostic biomarkers and therapeutic targets^38,39^. Among the detected spliceosome components, SF3B1 is a known HIV-1 dependency factor. SF3B1 is known to interact with the viral transactivator Tat, promoting RNA polymerase II recruitment to the HIV promoter, and is required for efficient viral transcription and latency reversal^40^. Pharmacologic or genetic inhibition of SF3B1 disrupts Tat-mediated transcription and prevents reactivation even in the presence of LRAs^40^. How changes in SF3B1 phosphorylation impact its functionality in the context of HIV latency reversal remains to be explored.

The activation of NF-κB is a canonical feature of latency reversal, which promotes nuclear translocation of transcription factors and recruitment of coactivators to the HIV long terminal repeat (LTR)^41,42^, which regulates proviral gene expression. The NF-κB signaling complex, represented by NFKB1, SKP1, and KPNA2, was identified in our data, including the activating NFKB1 S907 phosphorylation event^43,44^. The canonical NF-κB pathway is rapidly activated by inflammatory stimuli through IKKβ-mediated phosphorylation and degradation of IκBα, releasing p65/p50 to drive proinflammatory gene expression^45^. In contrast, the non-canonical pathway involves cytoplasmic sequestration of RelB by NFKB1 until its processing to p52, enabling p52/RelB nuclear translocation and sustained transcriptional programs^45^, including those associated with HIV-1 latency reversal^46^. SKP1 facilitates degradation of the inhibitory protein IκBα through the SCF ubiquitin ligase complex^47^, and KPNA2 mediates nuclear import of the NF-κB p65 subunit^48^. Interestingly, SKP1 and KPNA2 showed increased phosphorylation under SAHA treatment but decreased phosphorylation under PMA and prostratin, whereas NFKB1 exhibited the opposite pattern, with decreased phosphorylation in SAHA and increased phosphorylation in the PKC agonist conditions. These trends suggest that SAHA and PMA/prostratin may regulate NF-κB in opposing ways, which has been suggested previously^49^.

ERK was also present in our network and included MAPK1 (ERK2) and MAPK3 (ERK1). The ERK pathway is activated downstream of multiple receptor and kinase cascades, including PKC, which we observed in response to PMA and prostratin (Fig. 2F). ERK activity results in the phosphorylation of transcription factors and cofactors such as NF-κB, Sp1, and CREB, which enhance HIV-1 LTR activity^50–53^. In addition, we also found the Mediator complex, which bridges transcription factors to RNA polymerase II together with BRD4, a central chromatin-associated regulator of HIV transcription. BRD4 interacts with P-TEFb, the same transcriptional elongation complex recruited by the viral protein Tat to the HIV promoter^54^. In the absence of Tat, BRD4 can promote basal transcription by recruiting P-TEFb to the LTR. However, once Tat is expressed, BRD4 competes with Tat for P-TEFb binding, thereby restricting transcriptional elongation and helping maintain latency^55^. We observed a strong increase in BRD4 S601 in response to SAHA, with milder upregulation in response to PMA and prostatin, a site previously found to be upregulated in response to ultraviolet light^56^. The role of this site in regulating BRD4 binding to P-TEFb and/or HIV-1 transcription remains to be established.

Although less common, of the 39 complexes, 16 had at least one site that was significantly altered by all three LRAs, but not necessarily in the same direction (Fig. S2A). Complexes with the highest proportion of shared phosphosites included the ERK complex, TFTC, HES1 promoter-Notch enhancer, anti-HDAC2, ALL-1 supercomplex, and the CDC5L complex. These assemblies mediate transcriptional activation and chromatin remodeling, suggesting that distinct LRAs converge on phosphorylation of a shared nuclear regulatory layer that links signal transduction to gene expression.

Lastly, we integrated the core phosphoregulated sites with a published HIV-1 CRISPR screen by Hsieh et al. and identified 197 overlapping genes^57^, most of which localized to chromatin-related assemblies including the LCR-associated remodeling complex, CEN, WINAC, anti-HDAC2, SIN3-ING1b complex II, and the spliceosome (Fig. S2B). Together, our results demonstrate that LRAs rewire nuclear signaling through coordinated phosphorylation of protein complexes that regulate chromatin remodeling and RNA processing.

## DISCUSSION

Our study provides a comprehensive phosphoproteomics framework for understanding how LRAs remodel host signaling during HIV-1 reactivation. Across two independent experiments, we identified directionally consistent, abundance-independent phosphorylation sites, defining a reproducible and biologically coherent signature of LRA-induced signaling. Requiring reproducibility across experiments ensures that our core phosphoregulated sites represent coordinated cellular responses rather than stochastic variation. Furthermore, by excluding abundance-correlated sites, we specifically captured phosphorylation-driven regulation, uncovering signaling layers that accompany viral reactivation beyond changes in total protein abundance.

Network-level analysis showed that LRAs extensively remodel phosphorylation within nuclear protein complexes such as the spliceosome, Mediator, nuclear transport machinery, and NF-κB regulatory assemblies. The prominence of RNA processing and transcriptional complexes indicates that LRAs promote a signaling state that is conducive to reactivation of the provirus. Although SAHA, PMA, and prostratin act through distinct upstream mechanisms (HDAC inhibition versus PKC activation), they converge on a set of nuclear regulatory pathways and complexes that are known to regulate transcriptional elongation, chromatin accessibility, and RNA maturation. While PKC agonists are thought to drive viral transcription by activating NF-κB and HDAC inhibitors are thought to drive viral transcription through histone acetylation, this convergence suggests a conserved signaling architecture that may play a role in the transition from latency to reactivation, irrespective of the initiating LRA stimulus.

These findings have several therapeutic and mechanistic implications. Elucidating PTM-driven mechanisms of latency reversal could enable the rational design of targeted interventions to precisely modulate specific protein complexes or protein-protein interactions to promote reactivation within a “shock-and-kill” framework. Furthermore, the integration of our phosphorylation maps with lists of HIV-1 dependency factors or virus-host interactions could aid in the prioritization of host factors for further study.

While our analysis provides an extensive view of host phosphorylation remodeling, several limitations remain. The work was performed in a single J-Lat cell line model, which does not capture the full heterogeneity of the latent reservoir in people with HIV. Temporal resolution was also limited to 2 and 24 hour time points; intermediate and later stages may reveal additional dynamics in signaling and complex remodeling. Finally, phosphoproteomics alone cannot establish causal relationships between phosphorylation events and reactivation outcomes. Future studies aimed at directly mapping protein-protein interactions and identifying complexes regulated with LRA-induced phosphorylation changes will be essential to define the molecular hierarchy that drives these signaling networks.

## METHODS

### Culturing and reactivation J-Lat 10.6 cells

The J-Lat 10.6 cell line was from the NIH AIDS Reagent Program. The cells were maintained in RPMI-1640 medium (Gibco) supplemented with 10% fetal bovine serum (FBS, Gibco) and 1% penicillin/streptomycin (Gibco) at 37 °C and 5% CO₂ for at least one and a half week. Cells were passaged every 2-3 days at a 1:20 split ratio. For latency reversal experiments, 10 million J-Lat 10.6 cells were treated in a 10 cm cell culture dish for 24 hours under the following conditions: 0.01% DMSO (mock), 10 µM SAHA (Cayman Chemistry, Cat# 10009929), 5 ng/mL PMA (AdipoGen Life Sciences, Cat# AG-CN2-0010-M010), or 5 µM prostratin (Cayman Chemistry, Cat# 10272). Each condition has four technical replicates in each experiment. The collected cells were spun at 400 g for 5 min at room temperature to remove the cell culture media, and the pellet was washed two times with ice-cold 1× PBS (Corning).

### Quantifying reactivation via flow cytometry

A total of 50,000 cells per sample were transferred into an uncoated round-bottom 96-well plate (Corning, Cat# CLS3795). Cells were pelleted by centrifugation at 400 × g for 5 min at room temperature and washed once with 1× PBS (Corning, Cat# MT21040CV). After washing, cells were centrifuged again, and the supernatant was removed. The pellet was fixed in 200 μL of 1% paraformaldehyde (Electron Microscopy Sciences, Cat# 15714-S) for 15 min at room temperature. Following fixation, cells were centrifuged to remove paraformaldehyde, washed once with 100 μL of 1× PBS, and finally resuspended in 200 μL of 1× PBS. Flow cytometric analysis was performed on an Attune NxT Flow Cytometer (Thermo Fisher Scientific).

### Mass spectrometry sample preparation

Cells were lysed by adding 300 μL of lysis buffer (4% sodium deoxycholate, 0.1 M Tris-HCl, pH 8) and sonication using a tip sonicator (Fisher Scientific). The samples were then boiled at 95 °C for 5 min and centrifuged at 18,000 g for 10 min at 4 °C to clear out cell debris. The supernatant was transferred to a new protein LoBind (Eppendorf) tube, and protein concentration was estimated with the BCA assay kit (Pierce). Volumes corresponding to 1 mg of proteins were taken from each sample and normalized to 200 μL with the lysis buffer. Proteins were reduced with 10 mM TCEP (tris(2-carboxyethyl)phosphine, Pierce) at 56 °C for 30 min and alkylated with 40 mM 2-chloroacetamide (Thermo Scientific) at 23 °C in the dark for 30 min on a ThermoMixer (Eppendorf) at 800 rpm. The samples were diluted with 0.1 M Tris buffer (pH 8) to reduce the urea concentration to less than 1 M, and digested overnight at 23 °C with Trypsin (Promega) and Lys-C (Wako) at a protein:protease ratio of 1:100 (w/w). After digestion, Trifluoroacetic acid (TFA) was added to bring the pH to less than three. Peptides were desalted with the Oasis HLB 1 cc vac cartridge (Waters), and five percent of the eluted peptides were separated for global abundance proteomic analysis. The remainder was used for phosphopeptide enrichment. Both fractions were dried to completeness in a SpeedVac (Labconco). For phosphopeptide enrichment, MagReSyn Ti-IMAC HP beads (Resyn Biosciences) were aliquoted in a 1:2 (w/w) peptide:beads ratio and equilibrated three times with binding buffer (0.1 M glycolic acid in 80% acetonitrile (ACN), 5% TFA). Dried peptide samples were resuspended in 200 μL of binding buffer, added to the equilibrated beads, and incubated for 30 min at 23 °C, 1200 rpm. The unbound fraction was discarded, and the beads were washed sequentially with 200 μL each of the binding buffer, wash buffer 1 (60% ACN, 1% TFA, 200 mM NaCl), wash buffer 2 (60% ACN, 1% TFA), and finally with water (LC-MS grade, Fisher Scientific). Enriched phosphopeptides were eluted by incubating the beads with 150 μL of 1% (v/v) ammonium hydroxide (Sigma) in water (LC-MS-grade) for 10 min at 23 °C, 1200 rpm. The eluted peptides were transferred to a new protein LoBind tube (Eppendorf) containing 50 μL of 10% (v/v) formic acid (FA) in water (LC-MS grade). The elution step was repeated once more. Both eluates were pooled, dried in a SpeedVac (Labconco), and stored at -80 °C until further analysis.

### Mass spectrometry data acquisition

Dried peptides were resuspended in 0.1% (v/v) FA in water (LC-MS grade), and approximately 500 ng of peptides were analyzed on a timsTOF HT mass spectrometer (Bruker Daltonics), paired with a Vanquish Neo UHPLC system (Thermo Scientific). Mobile phase A consisted of 0.1% (v/v) FA in water (LC-MS grade), and mobile phase B consisted of 0.1% (v/v) FA in ACN (LC-MS grade). The LC was operated in trap-and-elute mode, where the peptides were first trapped onto a PepMap Neo Trap column (5 mm length, 100 Å pore size, 5 µm particle size) and then reversed-phase separated using gradients mentioned below on an Aurora Elite C18 reverse phase column (15 cm length, 75 µm I.D., 100 Å pore size, 1.5 µm particle size, IonOptiks) for Bruker CaptiveSpray source, kept at 50 °C using a column oven (Sonation Lab Solutions), and ionized in a CaptiveSpray source (Bruker Daltonics) at 1700 V. For global abundance proteomics analysis, the trapped peptides were separated on a gradient of mobile phase B as follows: 3% to 30% B over 51 min at 0.3 µL/min, then to 45% B in next 5 min, followed by an increase to 60% B over 1 min. The column was then flushed with 95% B for the next 3 minutes. For phosphoproteomics analysis, the same gradient was used, except the initial gradient was set 3% to 25% B over 51 min at 0.3 µL/min. For the global abundance analysis of 2 hours treatment conditions, the LC gradient was: 5% to 35% B over 37 min at 0.3 µL/min, then to 45% B in next 4 min, followed by an increase to 60% B over 1 min. The column was then flushed with 95% B for the next 3 minutes. For the phosphoproteomic analysis of 2 hours treatment conditions, the LC gradient was: 3% to 16% B over 22 min at 0.3 µL/min, then to 30% B in 15.5 min, and 45% B in next 4 min, followed by an increase to 60% B over 1 min. The column was then flushed with 95% B for the next 2.5 minutes. The dia-PASEF method for global proteomic and phosphoproteomic analysis was generated using the py_diAID^58^ software, with variable isolation window widths in the m/z vs ion mobility plane. The data were collected using 1 MS frame and 14 dia-PASEF frames per cycle, with an accumulation and ramp time of 100 ms, yielding an estimated mean cycle time of 1.59 s. Each dia-PASEF frame consisted of two ion mobility (1/K0) windows over the range 0.65-1.40 Vs cm-2. The mass range for dia-PASEF windows was 261.1-1350.8 m/z for global proteomic analysis and 274.8-1350.7 m/z for phosphoproteomics analysis. The collision energy was ramped linearly, from 59 eV at 1/K0 1.6 Vs cm-2 to 20 eV at 1/K0 0.6 Vs cm-2. For 2 hours treatment experiment (Experiment 3), the dia-PASEF methods for both global abundance proteomics and phosphoproteomics, were generated with timsControl Method Editor (ver. 7.02, Bruker Daltonics) software. For the global protein abundance analysis, dia-PASEF settings were: m/z range 250.2-1296.2, mobility range (1/K0) = 0.64-1.39 Vs cm−2, mass width = 20 Da, mass overlap = 1 Da, the number of MS/MS ramps = 14, and the number of MS/MS windows = 55. The estimated cycle time was 1.59s. The collision energy was ramped linearly, from 59 eV at 1/K0 1.39 Vs cm-2 to 20 eV at 1/K0 0.64 Vs cm-2. For phosphoproteomic analysis, dia-PASEF settings were: m/z range 279.8-1279.8, mobility range (1/K0) = 0.61-1.34 Vs cm−2, mass width = 28 Da, mass overlap = 1 Da, number of MS/MS ramps = 14, and the number of MS/MS windows = 37. The estimated cycle time was 1.59s. The collision energy was ramped linearly, from 59 eV at 1/K0 1.34 Vs cm-2 to 20 eV at 1/K0 0.61 Vs cm-2. The ramp and accumulation times for both dia-PASEF methods were set to 100 ms.

### Mass spectrometry data search and analysis

Raw files were searched against the Human UniProt SwissProt database (downloaded Aug. 31, 2024), appended with the HIV-1 J-Lat 10.6 proteome (downloaded May 25, 2024), using Spectronaut (ver. 19.5, Biognosys) directDIA workflow. For global proteome analysis, the default ‘BGS Factory Settings’ workflow was used with a few modifications. Briefly, the enzyme/cleavage rule was set to Trypsin/P (specific), with a maximum of two missed cleavages allowed per peptide. Acetylation (Protein N-term) and Oxidation (M) were set as variable modifications, and Carbamidomethylation (C) was set as a static modification. A maximum of five variable modifications was allowed per peptide. The false discovery rate (FDR) threshold for identification at the peptide spectrum match (PSM), peptide, and protein group levels was set to 1%. The precursor posterior error probability (PEP) and q value cutoffs were set to 0.2 and 0.01, respectively. The protein q value cutoff for the experiment and run was set to 0.01 and 0.05, respectively. For quantification, precursors that passed the q value threshold with the filtering setting ‘Identified (Qvalue)’ were further selected for MS2-level quantification by calculating the area under the curve between the XIC peak boundaries for each targeted ion. No imputation was performed, and cross-run normalization was enabled to apply label-free quantification within the experiment. The data were normalized using the ‘Global normalization’ strategy, which normalizes individual runs based on the median quantities of all peptides selected for normalization. For phosphoproteomics analysis, the default ‘BGS Phospho PTM’ workflow settings were used, with the following modifications: during Pulsar search, the PTM localization was set to True, and the minimum localization threshold was set to zero. For quantification, PTM localization was set to True, and the probability cut-off was set to zero to quantify all possible phosphorylation sites. Later, only those phosphosites with PTM site probability greater than 90% were used for all downstream analysis. Similar to the global proteome analysis, the normalization strategy was set to normalize individual runs to the global median peptide quantity. If not explicitly stated, all other parameters were set to the default Spectronaut settings. The Spectronaut candidates list containing unpaired t-test results, were exported for further downstream analyses in R (version 4.5.1). One PMA sample from experiment 2 of the phosphorylation dataset was excluded from further analyses due to poor correlation with its replicates (Fig. S1D). The global protein abundance and phosphoproteomic data analysis of 2 hours treatment conditions (Experiment 3) were conducted using Spectronaut (ver. 19.9, Biognosys) directDIA workflow with the same settings mentioned above.

### Gene set overrepresentation analysis

Gene set overrepresentation analysis (GSOA) was performed on lists of significant host genes based on the filtering steps for abundance and phosphoproteomics. GSOA uses the Gene Ontology Biological Processes and Cellular Component dataset from Molecular Signature Database (MSigDB). The background universe was defined as the HGNC names of all genes detected in our dataset. Pathway terms were ranked by the number of viruses associated with that term at an adjusted p-value below 0.05.

### Kinase activity analysis using prior knowledge networks

We performed kinase activity analysis on phosphoproteomics data, as done previously^59,60^. Log_2_ fold changes (Log2FC) in phosphosite intensities between treated and mock cells were calculated and used to infer kinase activity states using prior knowledge networks of kinase-substrate interactions derived from the Omnipath database^31^. Kinase activities were inferred as a z-score from a Z-test comparing Log2FCs in phosphosite measurements of the known substrates (per kinase) against the overall distribution of detected Log2FCs across the sample. This statistical approach has been previously shown to perform well at estimating kinase activities^32–34^.

### Proteomics data integration and filtering

Spectronaut output from two independent phosphoproteomics experiments (Exp1 and Exp2) was processed in R to identify reproducible and abundance-independent phosphorylation changes induced by LRAs. For each experiment, all PTM groups identified in the Spectronaut candidate lists (*i.e.*, Post Analysis tab) were imported and filtered to retain only entries corresponding to unique protein accessions. Rows containing multiple proteins (indicated by “;” in the ProteinGroups column) were removed to avoid ambiguity in protein assignment. Each phosphopeptide or protein abundance feature was annotated as significantly regulated if it met both thresholds: |Log₂FC| > 1 and q < 0.05. Phosphopeptide (PTM) and abundance (AB) data from each experiment were merged by their gene names and treatment conditions. Then, to distinguish phosphorylation-specific regulation from changes driven by total protein abundance, we implemented a filtering strategy based on three sequential criteria: 1) Identification of Commonly Detected Sites: Phosphorylation sites were first filtered to retain only those detected in both Exp1 and Exp2; 2) Directional consistency across experiments: Only phosphosites showing the same direction of phosphorylation and abundance regulation (up or down or not significant) in both Exp1 and Exp2 were retained; 3) Removal of Abundance-Dependent Phosphorylation Events: phosphosites exhibiting the same direction of change as their corresponding protein abundance (both upregulated or downregulated) were excluded, as these likely reflected abundance-driven effects. Sites were retained only when phosphorylation and abundance changes were opposite in direction (e.g., up-regulated phosphorylation on a down-regulated protein) or when both were not significant.

### Construction of protein complex networks

Protein interaction networks were generated in Cytoscape (v3.10.3)^61^ using phosphoproteins identified from the merged and abundance-corrected dataset. Only proteins showing significant phosphorylation changes in at least one condition were included. Protein-protein interactions and complex composition were derived from the CORUM database. Complexes containing fewer than four subunits were excluded to focus on stable, multi-protein assemblies. Each protein was assigned to the complex in which it has the most interactors, and network edges were restricted to interactions connecting retained complex members. To enrich for regulatory and signaling assemblies, only nuclear-localized proteins were kept based on subcellular localization annotations from the Human Protein Atlas^62,63^. The resulting nuclear CORUM network was visualized in Cytoscape. Donut rings representing individual phosphorylation sites were added using the Omics Visualizer plugin^64^, with segment colors corresponding to the log₂ fold change values for each site.

---

**Figure S1.**
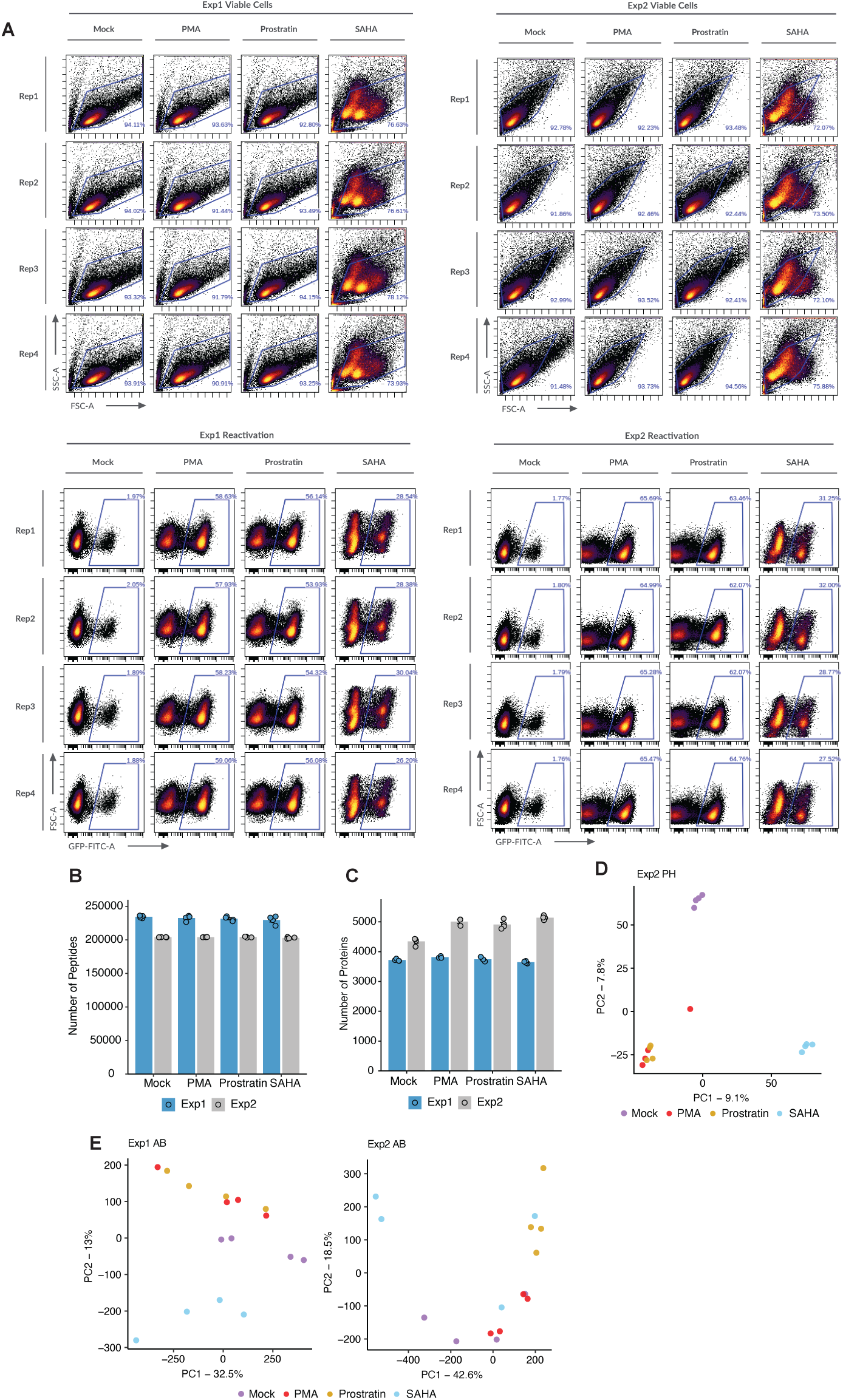
JLat flow cytometry and quality control results. (A) Flow cytometry FSC-A vs. SSC-A and FITC-A (GFP) vs. FSC-A graphs, with conditions as the columns and replicates as the rows. The gating shows viable cells and the GFP (reactivation) percentage. (B) Quantity of peptides detected in abundance proteomic data per LRA treatment and replicate across experiments 1 and 2. (C) Number of high confidence (Localization Probability > 0.9) phosphorylated proteins identified from the phosphoproteomics analysis per LRA treatment and replicate across Exp1 & 2. (D) PCA of experiment 2 phosphoproteomics(PH) data with all 16 samples, before removal of one of the PMA replicates. (E) PCA of abundance(AB) proteomics data from Exp1 & 2.

**Figure S2.**
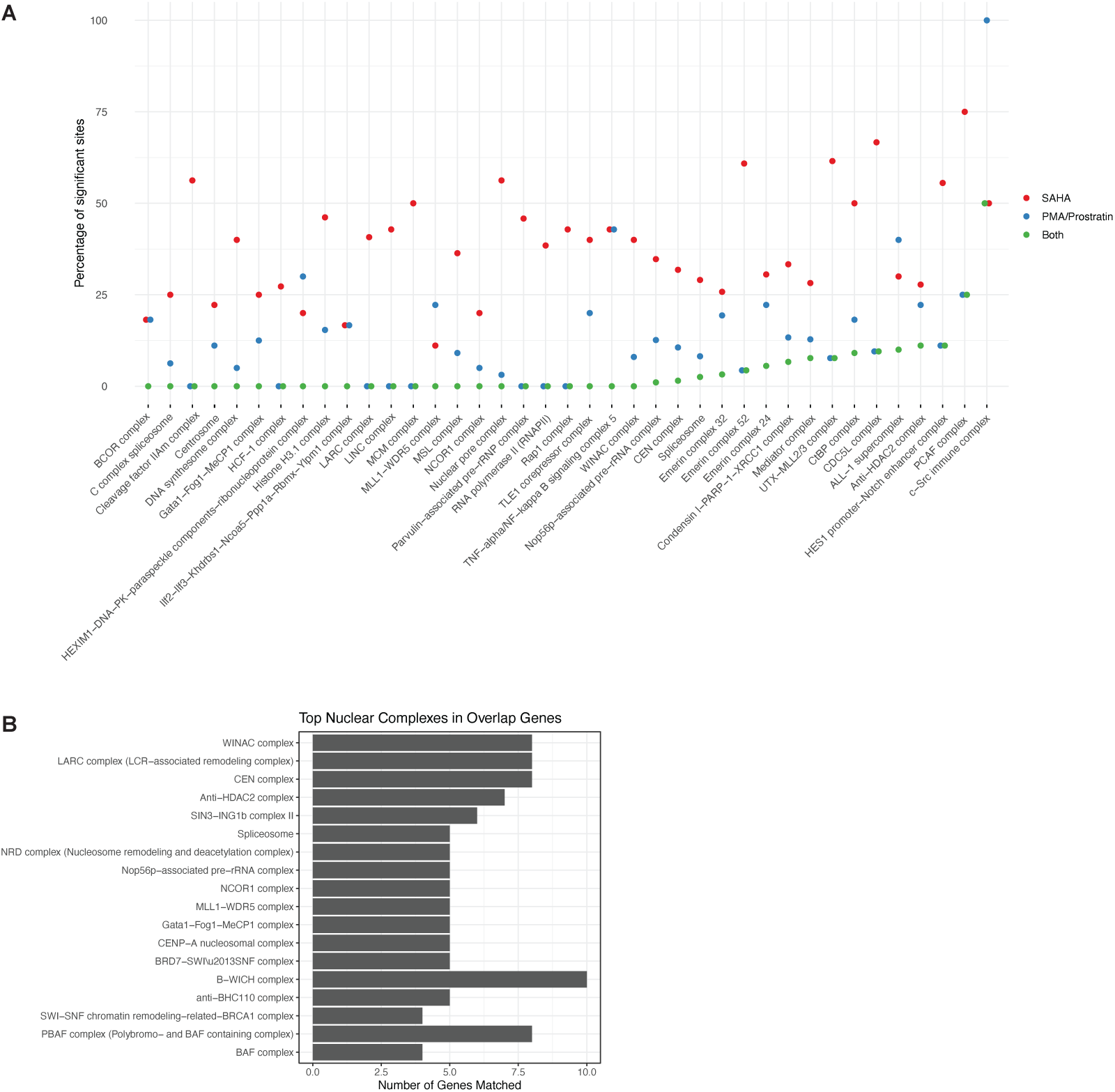
Overlap of phosphorylation sites between LRAs and with previously-reported knock-out screen dataset. (A) Comparison of phosphosite regulation across the 39 nuclear protein complexes affected by SAHA and PKC agonists (PMA and Prostratin). Complexes with overlapping phosphosites include c-Src signaling, TFTC, HES1 promoter–Notch enhancer, anti-HDAC2, ALL-1 supercomplex, and CDC5L. (B) Nuclear complexes involved in the overlap between LRA-regulated genes and a published HIV CRISPR screen by Hsieh et al..

## DATA AVAILABILITY

The supplementary data are provided as Excel tables. The raw mass spectrometry data will be deposited in the PRIDE database upon journal submission. Please contact the corresponding authors to request interim access.

## ACKNOWLEDGEMENTS

Our trainees were generously supported by the Howard Hughes Medical Institute (HHMI) Gilliam Fellowship (#GT17454 to SKM), Molecular Biology Institute Whitcome Fellowship to SKM, the UCLA-Caltech Medical Scientist Training Program (NIH-NIGMS T32GM008042 to SFB), the David Geffen Medical Student Scholarship to SFB, the Cellular and Molecular Biology T32 at UCLA (NIH-NIGMS T32GM145388 to DMW), the Microbial Pathogenesis Training Grant (MPTG NIH-NIAID T32AI007323 to YD), the Graduate Research Fellowship Program (GRFP) from the National Science Foundation (DGE-2444110 to YD), and the Undergraduate Research Scholars Program Grant funded by the Van Trees (USRP, 2025-2026 to JN).

## SUPPLEMENTARY TABLES

**Table S1. Unfiltered phosphorylation site changes.**

Spectronaut candidate output of all quantified post-translational modification from three independent datasets: two 24 hour LRA treatment experiments (Exp1 and Exp2) and one 2 hour treatment experiment (Exp3). Each entry includes the log₂ fold-change, q value, and significance status for SAHA, PMA, and Prostratin relative to mock control prior to any filtering.

**Table S2. Unfiltered total protein abundance changes.**

Spectronaut output of total protein abundance measurements from the same three experiments corresponding to Table S1. The table contains log₂ fold-change, q value, and significance classifications for all proteins detected under each treatment condition.

**Table S3. Abundance-corrected and merged phosphoproteomics dataset.**

Filtered phosphosite dataset obtained by merging the two 24 hour experiments (Exp1 and 2). Sites were retained only if they were consistently detected in both experiments, showed the same regulatory direction, and displayed phosphorylation changes independent of protein abundance. Phosphorylation and abundance log₂ fold-changes from the two experiments are included.

**Table S4. Gene set enrichment analysis summary.**

Results of enrichment analyses performed across multiple datasets. Includes (1) combined enrichment using both phosphorylation and abundance data across 24 hour and 2 hour experiments, (2) enrichment of final significant core phosphorylated proteins (Table S3) against Gene Ontology Biological Process (GOBP) terms, and (3) against Gene Ontology Cellular Component (GOCC) terms. Each sheet lists term identifiers, significance values, and associated site counts.

**Table S5. Kinase activity analysis summary.**

Predicted kinase activity scores in 24 hour treatment experiments. The table summarizes mean activity, z-scores, and statistical significance for kinases inferred to be up- or down-regulated by LRAs.

**Table S6. Nuclear complex phosphoprotein interaction network.**

Edge list representing the network of nuclear protein complexes affected by LRA treatment. Each edge connects a protein complex to its phosphorylated subunits or sites identified in the abundance-corrected phosphoproteome. This dataset and Table S3 underlie the network visualizations (Fig. 4) in the main text.

